# EEG decoding reveals neural predictions for naturalistic material behaviors

**DOI:** 10.1101/2023.02.15.528640

**Authors:** Daniel Kaiser, Rico Stecher, Katja Doerschner

**Author notes:** <u>Correspondence to:</u>.

## Abstract

Material properties like softness or stickiness determine how an object can be used. Based on our real-life experience, we form strong expectations about how objects should behave under force, given their typical material properties. Such expectations have been shown to modulate perceptual processes, but we currently do not know how expectation influences the temporal dynamics of the cortical visual analysis for objects and their materials. Here, we tracked the neural representations of expected and unexpected material behaviors using time-resolved EEG decoding in a violation-of-expectation paradigm, where objects fell to the ground and deformed in expected or unexpected ways. Participants were 25 men and women. Our study yielded three key results: First, both objects and materials were represented rapidly and in a temporally sustained fashion. Second, objects exhibiting unexpected material behaviors were more successfully decoded than objects exhibiting expected behaviors within 190ms after the impact, which might indicate additional processing demands when expectations are unmet. Third, general signals of expectation fulfillment that generalize across specific objects and materials were found within the first 150ms after the impact. Together, our results provide new insights into the temporal neural processing cascade that underlies the analysis of real-world material behaviors. They reveal a sequence of predictions, with cortical signals progressing from a general signature of expectation fulfillment towards increased processing of unexpected material behaviors.

**Significance Statement:** In the real world, we can make accurate predictions about how an object’s material shapes its behavior: For instance, we know that cups are typically made of porcelain and shatter when we accidentally drop them. Here, we use EEG to experimentally test how expectations about material behaviors impact neural processing. We showed our participants videos of objects that exhibited expected material behaviors (such as a glass shattering when falling to the ground) or unexpected material behaviors (such as a glass melting upon impact). Our results reveal a hierarchy of predictions in cortex: The visual system rapidly generates signals that index whether expectations about material behaviors are met. These signals are followed by increased processing of objects displaying unexpected material behaviors.

## Introduction

Many objects in our environment are made of a particular material, like porcelain, fabric, or rubber. Material properties critically determine how an object is used or interacted with. Thus, the ability to visually recognize material qualities quickly and correctly is important for planning actions and interactions (Buckingham et al., 2011, Klein et al., 2020).

Humans are able to make judgments about optical and non-optical material qualities, based on *visual* information alone (Adelson, 2001; Fleming, 2017; Kentridge & Chadwick, 2014; Paulun et al., 2019; Schmidt et al., 2017; Schmid & Doerschner, 2018; Schmid et al., 2021; van Assen et al., 2020). One possible explanation of this remarkable ability to infer non-optical material qualities, like softness or stickiness, from images is that we have learned over a lifetime to associate hand actions and haptic sensations with visual consequences of interactions (e.g., characteristic deformations).

This type of associative learning leads to strong expectations about how a material will behave under external forces (Alley et al., 2020; Bates et al., 2015; Paulun et al., 2017). For example, Bates et al. (2015) showed that humans can efficiently predict how liquids of different viscosities flow around solid obstacles. In our own work, we recently showed that existing expectations (i.e., those acquired through life-long learning) about the typical material properties of objects modulate perception (Alley et al., 2020; Malik et al., 2022). In our experiments, participants saw familiar objects (chairs, cups, custard) and unfamiliar novel shapes made of the same material as the familiar ones, fall to the ground. Upon impact the objects either behaved as expected (e.g., a cup shattering or a custard wobbling) or unexpectedly (e.g., a cup turning into liquid or wobbling). Only in the familiar object condition, we found that property ratings of the objects were systematically biased towards participants’ expectations about the material behavior and that unmet expectations were associated with longer response times that index additional processing demands. Together, these results demonstrate that the perception of real-world objects is invariably tied to expectations about their material behaviors.

We currently do not know how material behaviors are extracted across the neural visual processing cascade. More specifically, it is unclear at which stages of the processing cascade expectations about material behaviors modulate the cortical analysis of objects and their materials. To resolve these open questions, we devised an EEG experiment, in which we employed a variation of our previous paradigm where participants viewed real-world objects falling to the ground and exhibiting expected or unexpected material behaviors on impact. We then used time-resolved EEG decoding (Grootswagers et al., 2017) to track the representation of expected and unexpected material behaviors.

## Materials and Methods

### Participants

Twenty-five healthy adults (17 female, 8 male; mean age: 28.7 years, SD=7.4) participated in the experiment. All participants had normal or corrected-to-normal vision. Participants provided written informed consent before the experiment and received a monetary reimbursement. The study protocol was approved by the general ethical committee of Justus-Liebig-University Gießen. All experimental protocols were in accordance with the Declaration of Helsinki.

### Stimuli

Stimuli were eight unique full-color video renders (2s duration, 24Hz frame rate) depicting objects (chair, milk, custard, glass) falling from a fixed starting point down to the ground (Fig. 1a). Upon hitting the ground, an object either displayed its expected (e.g., custard wobbling on impact) or an unexpected (e.g., custard shattering to pieces on impact) material behavior (Fig. 1b). To create expected and unexpected stimuli, material behaviors were swapped among two pairs of objects: (1) the chair could stay rigid or splash like a liquid and the milk could splash or become rigid, and (2) the custard could wobble or shatter to pieces and the glass could shatter to pieces or wobble (Fig. 1b). There were thus two material behaviors associated with each object. Each resulting combination of object and material behavior was conveyed through a single unique video. The videos were used in previous behavioral studies on material perception and are available at https://doi.org/10.5281/zenodo.2542577.

**Figure 1.**
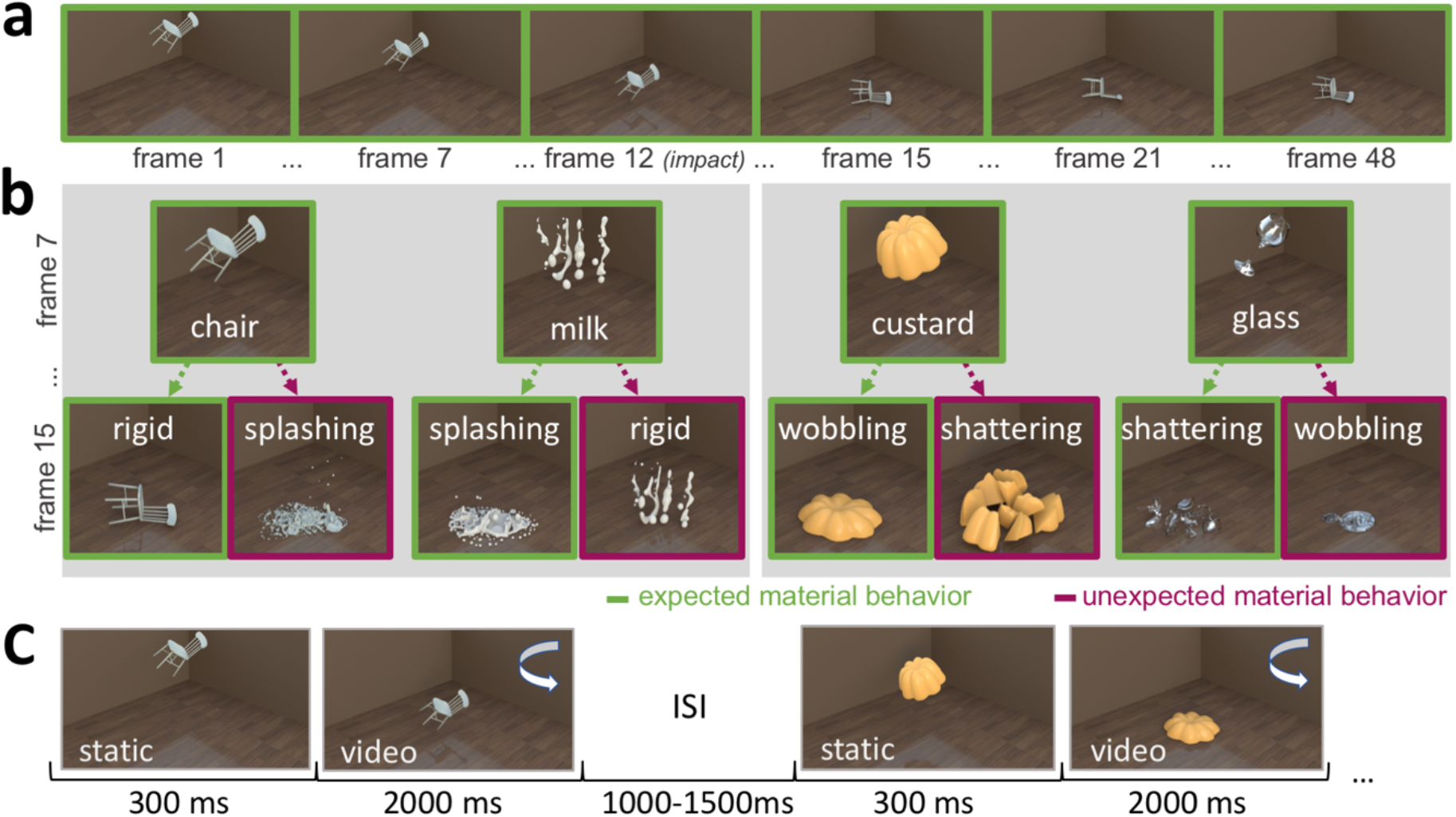
Stimuli and paradigm. (**a)** Example movie sequence for the object chair, subsampled at different frames throughout the video. (**b)** Illustration of the different object-material behavior combinations in our experiment at two time points during the animation (frames 7 and 15). Green color denotes expected material behaviors, e.g., the chair falling to the ground rigidly, purple color denotes unexpected material behaviors, e.g., the chair splashes upon hitting the ground. To create the unexpected material behaviors, two objects “swapped” material behaviors between them: chair - milk and custard - glass. Stimuli can be downloaded at https://doi.org/10.5281/zenodo.2542577. (**c)** Experimental paradigm. Participants viewed the videos in a random sequence while detecting occasional luminance dims in the whole stimulus.

The objects were rendered at approximately the same size so that they would behave in a similar way under gravity and were matched for motion energy across the expected and unexpected material behaviors (see Alley at al., 2020, for a more detailed description of the stimuli).

### Paradigm

Stimulus presentation was controlled using the Psychtoolbox (Brainard, 1997) for Matlab. Stimuli were presented on an AORUS FI32Q monitor at a refresh rate of 170Hz. The whole videos subtended 21 degrees by 16 degrees visual angle. Each trial (Fig. 1c) started with a 300ms static presentation of the first video frame. After that, the remaining video was played (47 frames with a frame time of 41.2ms each), with the object’s material behavior becoming apparent upon impact after 712ms relative to the first onset of the static presentation (i.e., on the 12th video frame). The impact frame was identical across all videos. Participants were instructed to maintain central gaze and to refrain from blinking during the stimulus presentation. They were further asked to respond to slight, but noticeable luminance dims in the videos by pressing the space bar. These luminance dims could only occur between the 13th and 36th frame, with onsets drawn from a truncated normal distribution peaking at the 33rd frame (i.e., 1,577ms after onset). We made targets more likely to appear late in the trial to ensure sustained attention during the trial. Participants on average detected the luminance dims in 80% of the cases (due to technical problems, responses were only recorded for 16 participants). Target trials were not used in the EEG analyses. Trials were separated by an inter-trial interval randomly varying between 1,000 and 1,500ms. The whole experiment featured 960 trials (including 96 target trials), and each video was presented equally often. Trial order was fully randomized.

### EEG recording and preprocessing

EEG signals were recorded using an Easycap 64-channel system and a Brainproducts amplifier, at a sample rate of 500Hz. Electrodes were arranged according to the standard 10-10 system. Data preprocessing was performed using the FieldTrip toolbox (Oostenveld et al., 2011) for Matlab and followed our previously established preprocessing routines for multivariate EEG decoding studies (Kaiser, 2022; Kaiser et al. 2019, 2020). The data were epoched from -500ms to 2800ms relative to stimulus onset, band-stop filtered to remove 50Hz line noise, re-referenced to the average across all electrodes, downsampled to 200Hz, and baseline corrected by subtracting the mean pre-stimulus signal. No high- or low-pass filters were applied to avoid spurious temporal distortions in the filtered data (van Driel et al., 2021; VanRullen, 2011). After that, noisy channels were removed by visual inspection. Eye artifacts were removed using independent component analysis (ICA) and visual inspection of the resulting components. No trials were removed during preprocessing. Finally, the data were further downsampled to 100Hz before analysis, to increase signal-to-noise-ratio while decreasing the number of individual analysis time point in our relatively long analysis epochs.

### Decoding analyses

Decoding analysis was performed using the CoSMoMVPA toolbox (Oosterhof et al., 2016) for Matlab, and carried out separately for each participant. All analysis were performed in a time-resolved fashion (Grootswagers et al., 2017), that is, separate analyses were conducted at each time point (i.e., for steps of 10ms). Classifiers were trained and tested on voltage patterns across all available EEG electrodes. Specifically, linear discriminant analysis (LDA) classifiers were always trained on one subset of the data and tested on a disjoint subset of the data (see below for details on the different cross-validation procedures). Classifier accuracies were averaged across all possible train-test splits to yield a time course of decoding accuracies. All decoding time courses were smoothed with a 3 time-point (i.e., 30ms) moving average (Kaiser et al., 2016a). Statistical testing was then performed across participants (see below for details). We performed multiple decoding schemes to retrieve complementary stimulus attributes, which are detailed in the following (also see Table 1 for an overview).

**Table 1.** Summary of all decoding schemes used in the study.

| Decoding Scheme | Example Training Conditions | Example Testing Conditions | Interpretation | Reported in Figure |
| --- | --- | --- | --- | --- |
| **Object decoding (uncontrolled)** | Glass shattering (1) vs. Custard wobbling (2) | Glass shattering (1) vs. Custard wobbling (2) | Object information (potentially confounded by material) | 2a |
| **Object Decoding (across materials)** | Glass shattering (1) vs. Custard wobbling (2) | Glass wobbling (1) vs. Custard shattering (2) | Object information (independent of material) | 2a |
| **Material Behavior Decoding (uncontrolled)** | Glass shattering (1) vs. Custard wobbling (2) | Glass shattering (1) vs. Custard wobbling (2) | Material information (potentially confounded by object) | 2b |
| **Material Behavior Decoding (across objects)** | Glass shattering (1) vs. Custard wobbling (2) | Custard shattering (1) vs. Glass wobbling (2) | Object information (independent of material) | 2b |
| **Expected Material Behavior Decoding** | Glass shattering (1) vs. Custard wobbling (2) | Glass shattering (1) vs. Custard wobbling (2) | Information about expected combinations (potentially confounded by object and material)* | 3 |
| **Unexpected Material Behavior Decoding** | Glass wobbling (1) vs. Custard shattering (2) | Glass wobbling (1) vs. Custard shattering (2) | Information about unexpected combinations (potentially confounded by object and material)* | 3 |
| **Expectation Decoding (uncontrolled)** | Glass wobbling (1) vs. Milk melting (2) | Glass wobbling (1) vs. Milk melting (2) | Information about expectations (potentially confounded by object and material) | 4 |
| **Expectation Decoding (across objects and materials)** | Glass wobbling (1) vs. Milk melting (2) | Chair melting (1) vs. Custard Wobbling (2) | Information about expectations (independent of object and material) | 4 |

*\*Note that both expected and unexpected material behavior decoding are similarly confounded by object and material, allowing for a fair comparison between the two decoding schemes.*

#### Object decoding (uncontrolled)

Each object (chair, milk, custard, glass) produced data from expected and unexpected material behavior trials. Here, our goal was to decode between the different objects shown on each trial, irrespective of their material behavior. This was done in a 10-fold cross-validation scheme: we assigned 90% of trials for each unique video to the training set and the remaining 10% of trials to the testing set. Each individual video thus appeared both in the training and testing set. During training, different labels were assigned to videos containing the four objects, and during testing classifiers had to predict the correct object label. The classification procedure was repeated 10 times, until each chunk served as the test set once; accuracies were averaged across these 10 repetitions. Performing the classification in this manner allows for the possibility that the classifier not necessarily picks up on object identity per se, but instead on idiosyncratic aspects of the movies, e.g., the specific material behaviors of each object.

#### Object decoding (across material behaviors)

As the uncontrolled object decoding scheme may lead to an overestimation of object information (because classifiers can pick up on material behaviors common between the train and test sets), we performed a second analysis that abolished such commonalities between the train and test sets. Specifically, classifiers were now always trained on objects exhibiting a different material behavior than they exhibited during testing (e.g., training on a glass shattering, but testing on a glass wobbling), thus only allowing them to capitalize on the object identity and not the material behavior. Results were averaged across 16 possible assignments of these halves and across both train-test directions (e.g., train: chair rigid; test: chair liquid and vice versa, for each of the four objects).

#### Material behavior decoding

Like the object decoding, material behavior decoding was assessed in two analyses. First, we performed a 10-fold cross-validation analysis, where the same videos appeared in the train and test sets (uncontrolled analysis). Second, material behavior decoding was assessed across objects. Here, we trained classifiers on material behaviors exhibited by one object and tested them on the same material behaviors exhibited by a different object (e.g., training on a glass shattering, but testing on a custard shattering). Analysis details were identical to the object decoding described above.

#### Decoding for objects exhibiting expected and unexpected material behaviors

Here, we performed two separate decoding analyses, which separately tracked the representations for expected and unexpected object-material combinations. For each analysis, we only used the data for objects displaying expected or unexpected material behaviors, respectively. Data were split into 10 equally sized chunks: we assigned 90% of trials for each unique video to the training set and the remaining 10% of trials to the testing set. Each individual video thus appeared both in the training and testing set. During training, different labels were assigned to the four different videos, and during testing classifiers had to predict the correct video. The classification procedure was repeated 10 times, until each chunk served as the test set once; accuracies were averaged across these 10 repetitions. By comparing the decoding timeseries for the expected and unexpected material behaviors, we could infer whether decoding is enhanced for unexpected material behaviors, where unpredictability may require additional visual processing.

#### Expectation decoding

Here, we directly decoded between objects that displayed an expected or an unexpected material behavior. First, this was done by splitting the data into 10 equally sized chunks: we assigned 90% of trials for each unique video to the training set and the remaining 10% of trials to the testing set. Each individual video thus appeared both in the training and testing set. During training, two different labels were assigned to videos, reflecting whether the material behavior was expected or unexpected, and during testing classifiers had to predict the correct expectation label. The classification procedure was repeated 10 times, until each chunk served as the test set once; accuracies were averaged across these 10 repetitions. Second, to abstract away from specific objects and materials shown on individual trials, we trained classifiers on the first pair of object and material behaviors (chair/milk – rigid/liquid) and tested classifiers on the second pair of objects and material behaviors (custard/glass – wobbling/shattering), or vice versa. As classifiers encountered different objects and material behaviors in the training and test sets, successful decoding in this analysis reveals a general signal of expectation fulfillment. Results were averaged across both train-test directions.

### Statistical testing

Decoding accuracies were tested against chance level using one-sided t-tests, separately across time. For comparing decoding accuracies, two-sided tests were used. P-values were corrected for multiple comparisons across time using false-discovery-rate (FDR) corrections. Only tests after stimulus onset and tests yielding at least two consecutive timepoints reaching statistical significance were considered. T-statistics and Cohen’s d as a measure of effect size are reported for all peak effects.

### Data availability

Stimuli are available on at https://doi.org/10.5281/zenodo.2542577. Data are available at https://osf.io/2bqav. Other materials are available on request.

## Results

To track the emergence of neural representations of objects and material behaviors, we used time-resolved multivariate decoding analyses. These analyses yielded a time course of when objects and materials are discriminable from EEG sensor patterns, as well as when representations are influenced by expectations about material behaviors.

### Object decoding

To track object representations across time, we first trained classifiers on discriminating videos that contained different objects, irrespective of the material behavior (Fig. 2a). In this uncontrolled decoding scheme, classifiers had access to all available videos in both the train and test sets. These classifiers could successfully predict the objects from 90ms and across the whole epoch (peak at 720ms, t[24]=8.3, d=1.7). This emergence of object decoding is consistent with decoding of objects in static images (Contini et al., 2017). As classifiers in this uncontrolled decoding scheme are trained and tested on the same trials, they can capitalize on idiosyncrasies in individual stimuli, which include the material behavior. We thus performed a second analysis where classifiers were trained on discriminating the objects for one set of material behaviors and tested on the same objects exhibiting different material behaviors. In this analysis, objects were successfully decoded from 100ms to 950ms (peak at 720ms, t[24]=7.3, d=1.5). It is worth noting that each object in our study is only conveyed through two videos – which are identical up to the impact frame – so that object information until the impact encompasses all low- and mid-level features that define an individual object. After the impact, however, object representations did not persist for more than 300ms, suggesting that the idiosyncratic material behaviors abolished persistent object information.

**Figure 2.**
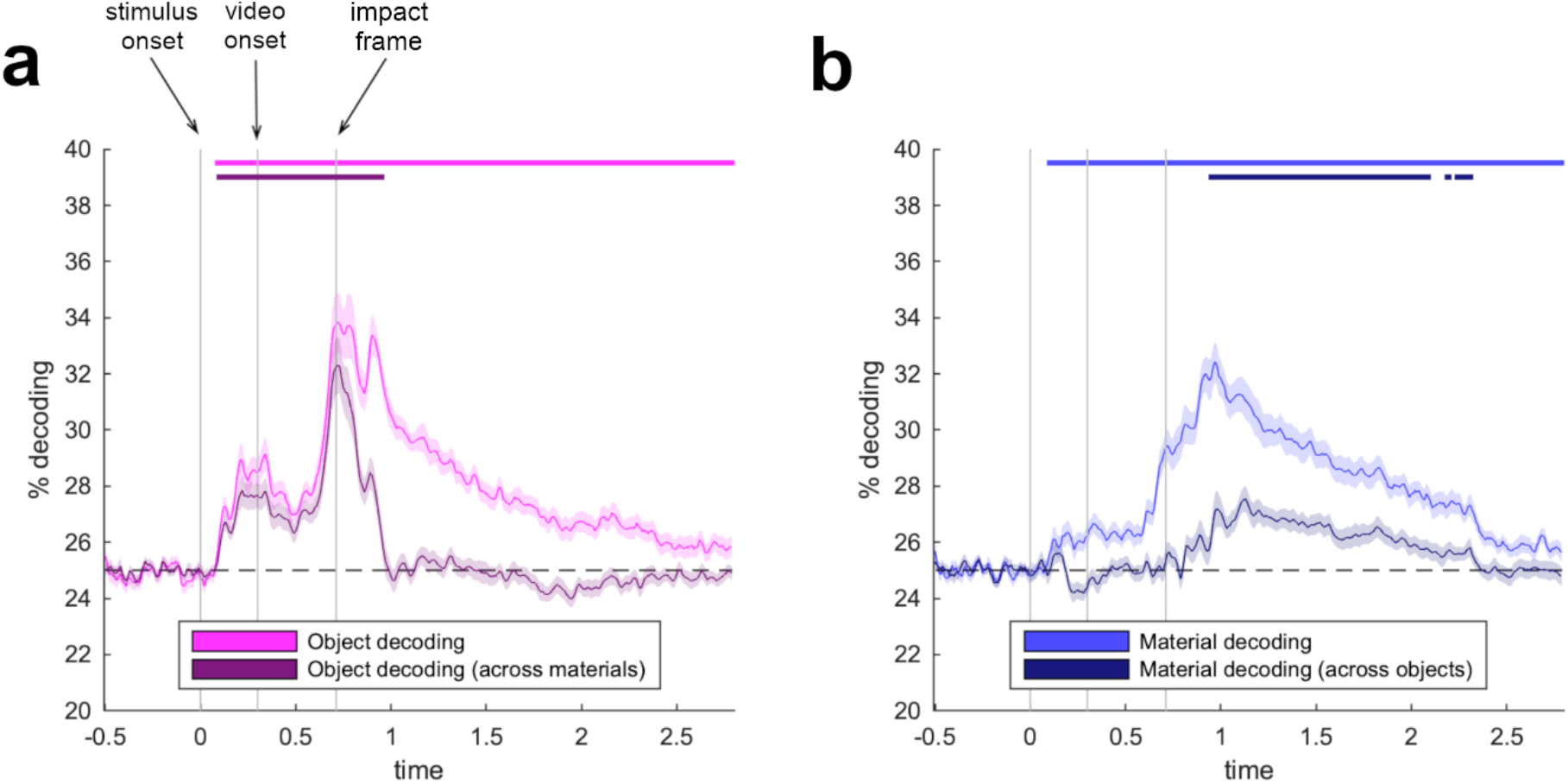
Neural representations of objects and materials across time. Both objects **(a)** and materials **(b)** were reliably decoded from EEG signals. When classifiers needed to generalize across the different material behaviors, object decoding vanishes about 300ms after the object hits the ground (impact frame). When classifiers needed to generalize across the different objects, material decoding emerged 240ms after the impact frame, which is when the material behavior is revealed. Error margins represent standard errors of the mean. Significance markers denote p<0.05 (corrected for multiple comparisons across time).

### Material behavior decoding

To track material representations across time, we first trained classifiers on discriminating videos that contained different material behaviors, irrespective of the objects that exhibited these behaviors (Fig. 2b), again using all available videos in the train and test sets. These classifiers could successfully predict the material behaviors from 100ms and across the whole epoch (peak at 970ms, t[24]=10.2, d=2.0). As classifiers in this uncontrolled decoding scheme are again trained and tested on the same trials and could thus also pick up on object information, we performed a second analysis where classifiers were trained on discriminating the material behaviors for one set of objects and tested on the same material behaviors exhibited by a different set of objects. In this analysis, material behaviors were successfully decoded from 950ms to 2,310ms (peak at 1,130ms, t[24]=5.5, d=1.1), providing evidence for a neural representation of material behavior that is formed around 240ms after the object hits the ground.

### Decoding for objects exhibiting expected and unexpected material behaviors

We next asked whether objects displaying expected and unexpected material behaviors give rise to cortical representations of different qualities: that is, are objects’ unexpected material behaviors better discriminable, because unmet predictions lead to recurrent (Urgen & Boyaci, 2021) or enhanced processing of the visual input? To answer this question, we trained two separate classifiers on discriminating videos that contained an expected material behavior and on discriminating videos that contained an unexpected material behavior, respectively (Fig. 3). Both classifiers successfully discriminated between the videos, for objects exhibiting expected material behaviors from 130ms to 2,710ms (peak at 2,310ms, t[24]=7.4, d=1.5) and for objects exhibiting unexpected material behaviors from 90ms to the end of the epoch (peak at 920ms, t[24]=12.1, d=2.4). Both of these individual analysis – given that the same videos were used in the train and test sets – conflate information about the videos’ visual properties, the object, and the material. However, they both do so in a comparable way, so that the difference in decoding for expected and unexpected material behaviors can be fairly assessed. Critically, we found enhanced decoding for the objects exhibiting unexpected material behaviors, compared to those displaying expected behaviors, from 900ms to 1,000ms (peak at 920ms, t[24]=5.1, d=1.0). This shows that already around 190ms after the material behavior is revealed (i.e., after the object hits the ground), there is a boost in cortical representations for objects displaying unexpected material behaviors. This enhancement may reflect additional processing demands when predictions about material behaviors are not met. The enhanced processing can in principle result from an enhanced coding of the object (following an unexpected material behavior), an enhanced coding of the material behavior (exhibited by an unlikely object), or by both at the same time.

**Figure 3.**
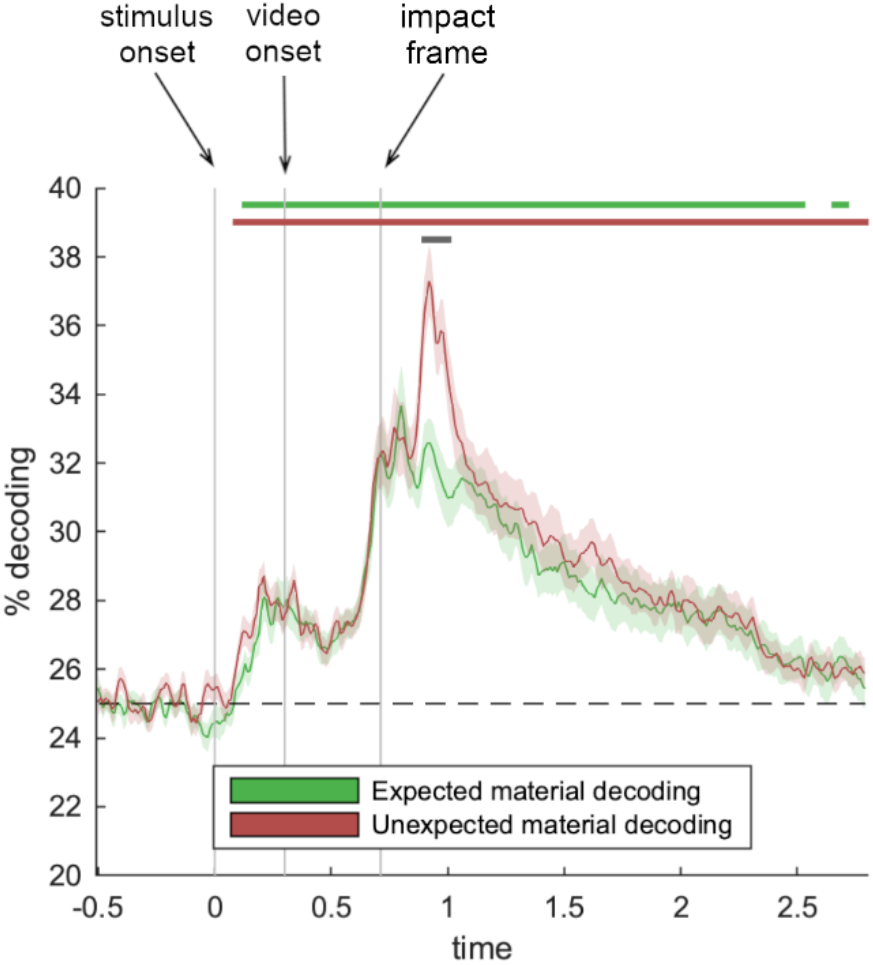
Enhanced representation of unexpected materials. When comparing decoding for objects that displayed expected and unexpected material behaviors, we found enhanced representations of objects exhibiting unexpected material behaviors that occurred around 190ms after the material was revealed (grey significance markers). Error margins represent standard errors of the mean. Significance markers denote p<0.05 (corrected for multiple comparisons across time).

### Expectation decoding

Finally, we asked whether there is a more general neural signal that indexes violations of material expectations that generalize across different objects and material behaviors. Such a signal would indicate that there is a generic implementation of a prediction error that either triggers subsequent differences in visual processing or that, alternatively, follows from such differences. To answer this question, we trained classifiers on discriminating between all videos that contained an expected material behavior and all videos that contained an unexpected material behavior, in a two-way classification analysis (Fig. 4). In a first analysis, this way again done in an uncontrolled decoding scheme, where the same videos were present in the train and test sets. Here, classifiers successfully discriminated between expected an unexpected videos from 220ms to the end of the epoch (peak at 1,020ms, t[24]=7.3, d=1.5). However, as these classifiers are again trained and tested on identical videos and thus can capitalize on pixel similarities in the train and test videos, we probed neural signals related to the fulfillment of expectations in a second analysis: Here, we trained classifiers on one combination of objects and materials (chair/milk – rigid/liquid) and tested them on another combination of objects and materials (custard/glass – wobbling/shattering), so that the classifiers could neither learn information about specific objects nor information about specific materials. In this analysis, we also found significant decoding of material expectations at multiple time points between 820ms and 1,300ms (peak at 1,130ms, t[24]=4.2, d=0.8). Interestingly, the first expectation-related decoding therefore occurred within 150ms after the impact frame, preceding the enhanced representation of unexpected material behaviors reported above. This pattern of results unveils a cascade of neural events, where an initial prediction error signal reflects a violation of expectation (indexing that the input does not match the expected visual pattern). This initial signal then triggers enhanced visual processing of unexpected material behaviors.

**Figure 4.**
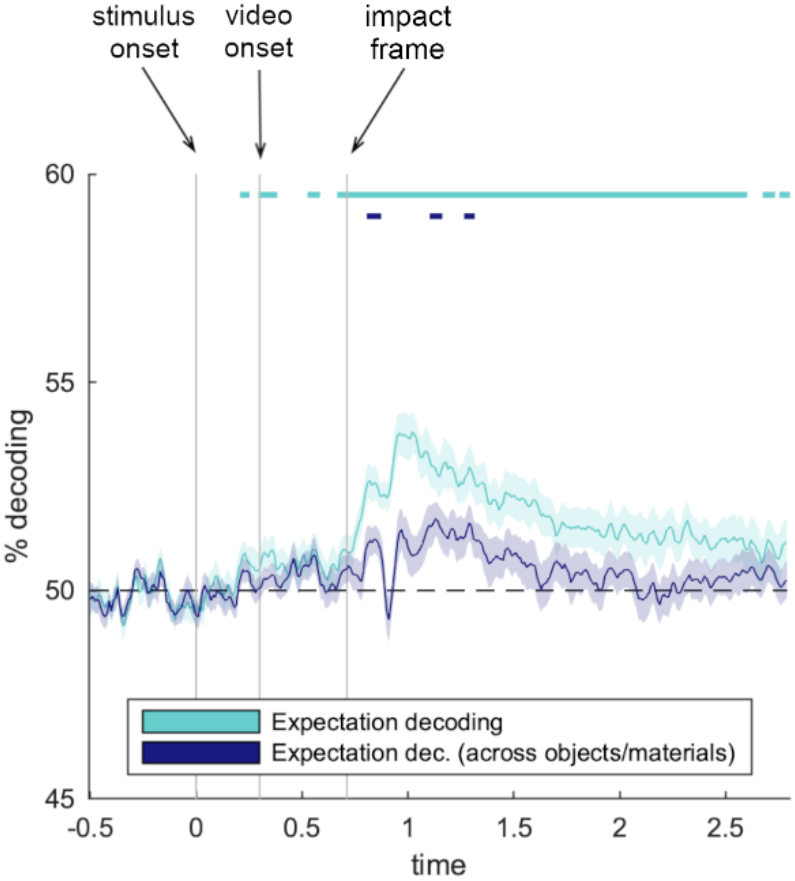
A general neural signal of expectation for material behaviors. In a two way-decoding, we found that neural signals contained reliable information about whether the material behavior was expected or unexpected. Critically, when classifiers were trained and tested across different objects and materials, we still found an early, general signal reflecting participants’ expectation for objects’ material behaviors, which occurred within 150ms of the object hitting the ground. Error margins represent standard errors of the mean. Significance markers denote p<0.05 (corrected for multiple comparisons across time).

## Discussion

Here, we used time-resolved EEG decoding to reveal the representation of expected and unexpected material behaviors in real-world objects. Our study yielded three key results: First, both objects and materials are represented in a temporally sustained fashion. Second, expected materials behaviors of objects are represented more strongly than unexpected behaviors within 190ms after the impact. Third, general signals of expectation fulfillment that generalize across specific objects and materials are found within the first 150ms after the material is revealed. Together, our results provide new insights into the temporal neural processing cascade that underlies expectations for real-world object behaviors: When material behaviors are expected from an object’s typical behavior in the world, the material is not strongly encoded in neural signals (indexed by decoding performance decreasing over time after the material is revealed). By contrast, when material behaviors are unexpected, material representations are enhanced (indexed by a boost in decoding performance after the material is revealed), reflecting the need for further visual analysis when the material behavior cannot be anticipated from the outset. This enhancement may be triggered by a general expectation-related signal that emerges rapidly, within 150ms after the material becomes apparent.

The early, general signal indexing fulfillment versus violation of expectation for material behaviors can be conceived as a neural prediction error (Rao & Ballard, 1999; Friston, 2005, 2010), where unmet expectations trigger error signals. The timing of this signal is consistent with prediction errors for expected simple stimuli and objects (Johnston et al., 2017; Robinson et al., 2018; Stefanics et al., 2015; Tang et al., 2018). Interestingly, Hogendoorn & Burkitt (2018) reported that neural signals at around 150ms post-stimulus signal the fulfillment of expectations about object movement trajectories. The early expectation-related signals observed in our study may reflect a similar mechanism, as an object’s material was conveyed through different movement patterns when hitting the ground. At this stage of processing, prediction errors in material perception may be triggered by predicted movement patterns: Already before impact, the brain may form expectations about the concerted transformation of low- and mid-level features on impact, which in turn leads to prediction errors that are similar across the individual objects exhibiting unexpected behaviors. The general signal of expectation fulfillment found here peaks first after 150ms from the impact, but a second peak emerges shortly after 400ms from the impact. In our study, this late peak is indeed statistically more robust than the early 150ms peak. Future studies are thus needed to solidify this temporal pattern of general expectation signals and to clarify the different functional roles of the early and late components of these signals.

We show that this general signaling of fulfilled expectations is followed by stronger representation of objects exhibiting unexpected material behaviors from 190ms within the impact. In principle, this effect can be explained in two ways: First, there may be an increase in processing for objects displaying unexpected material behaviors. Models of Bayesian inference (Kersten et al., 2004) predict that unmet expectations lead to recurrent updating of priors which requires additional processing of objects that exhibit unexpected material behaviors. Such recurrent prior updating can also explain delayed behavioral responses to objects displaying unexpected material behaviors (Urgen & Boyaci, 2022). Second, there may also be a decreased need for processing objects that exhibit expected material behaviors. When the material behavior can be expected, predictions can efficiently “explain away” the sensory input and further processing is eased (Friston, 2005, 2010). Although both mechanisms may play out concurrently, future studies could additionally use stimuli for which no expectations about material behavior are formed (such as meaningless shapes) to effectively dissociate the two mechanisms.

More generally, our study also provides new insights into the timing of material representations in the brain. In our study, robust material representations formed from 240ms after impact. This is considerably later than the representations observed in another EEG study on material perception from texture (Wiebel et al., 2014). This difference in timing may reflect differences between material perception from textures and movement. However, the later timing in our study is also attributable to our analysis scheme, in which we carefully probed material representations that generalize across objects, as well as across expected and unexpected material behaviors. It is worth noting that material representations may occur earlier than reported here when they are not probed across expected and unexpected cases.

The exact timing of material representation requires additional studies. In the absence of such studies in the domain of material perception, the timing of our effects can be contextualized with respect to EEG studies in the domain of object and scene perception. Several studies have argued that basic object perception, such as the extraction of object category (Carlson et al., 2013; Cichy et al., 2014; Contini et al., 2017; Kaiser et al., 2016b), is accomplished early, within the 200ms of processing. However, other work suggests that more complex perceptual analysis, such as the extraction of high-level scene attributes (Cichy et al., 2017; Harel et al., 2016; Kaiser et al., 2020) or the processing of objects embedded in scenes (Graumann et al., 2022) occurs after 200ms. At this time, representations may however already be altered by feedback from cognitive processes. Whether a latency of 240ms for extracting material properties can be strictly attributed to perceptual analysis is therefore still unclear.

The neural timing of material representation can also be related to behavioral response times. Judgements of materials (e.g., “Is this plastic?” or “Is it warm or cold?”) take about 100ms longer (median 532ms) than judgements of simple visual features such as color (434ms) or orientation (426ms; Sharan et al., 2014), mirroring the delay in neural decoding of objects versus materials. In a recent behavioral study, observers made two-alternative-forced-choice judgments (“Did it break?”) about materials that, similar to this study, either behaved expectedly or surprising (Malik et al., in press). Here, response times in the expected condition where comparable to that found by Sharan et al. (2014) – on average 510ms – and about 100ms longer in the surprising condition. Together, these behavioral results suggests that material processing is rapid, but that it also varies with the expectations and task demands.

In sum, we show that the neural representation of material behaviors is tightly linked to the expectations we form based on our real-world experience. Expected and unexpected material behaviors lead to differing representations across the visual processing cascade, with early signals reflecting a general signature of expectation fulfilment and later signals reflecting increased processing of unexpected, compared to expected material behaviors. The emergence of both these effects within the first 200ms of processing suggests that material representations are formed at fundamental stages of perceptual analysis.

## Acknowledgements

D.K. and K.D. are supported by the Deutsche Forschungsgemeinschaft (DFG SFR/TRR 135). D.K. is also supported by an ERC Starting Grant (ERC-2022-STG 101076057). This research was further supported by “The Adaptive Mind”, funded by the Excellence Program of the Hessian Ministry of Higher Education, Science, Research and Art. Thanks to Marius Geiss for help with EEG data collection.

## References

Adelson, E. H. (2001). On seeing stuff: The perception of materials by humans and machines. In Rogowitz B. E. &Pappas T. N. (Eds.), Proceedings of SPIE: Vol. 4299. Human Vision and Electronic Imaging VI, 1–12.

Alley, L. M., Schmid, A. C., & Doerschner, K. (2020). Expectations affect the perception of material properties. Journal of Vision, 20(12), 1–1.

Bates, C., Battaglia, P. W., Yildirim, I., & Tenenbaum, J. B. (2015, July). Humans predict liquid dynamics using probabilistic simulation. In CogSci.

Brainard, D. H., (1997). The psychophysics toolbox. Spatial Vision, 10(4), 433–436.

Buckingham, G., Ranger, N.S. & Goodale, M.A. The material–weight illusion induced by expectations alone. Atten Percept Psychophys 73, 36–41.

Carlson, T., Tovar, D. A., Alink, A., & Kriegeskorte, N. (2013). Representational dynamics of object vision: the first 1000 ms. Journal of vision, 13(10), 1–1.

Cichy, R. M., Khosla, A., Pantazis, D., & Oliva, A. (2017). Dynamics of scene representations in the human brain revealed by magnetoencephalography and deep neural networks. NeuroImage, 153, 346–358.

Cichy, R. M., Pantazis, D., & Oliva, A. (2014). Resolving human object recognition in space and time. Nature Neuroscience, 17(3), 455–462.

Contini, E. W., Wardle, S. G., & Carlson, T. A. (2017). Decoding the time-course of object recognition in the human brain: From visual features to categorical decisions. Neuropsychologia, 105, 165–176.

Fleming, R. W. (2017). Material perception. Annual Review of Vision Science, 3, 365–388.

Friston, K. (2005). A theory of cortical responses. Philosophical Transactions of the Royal Society B: Biological Sciences, 360(1456), 815–836.

Friston, K. (2010). The free-energy principle: a unified brain theory? Nature Reviews Neuroscience, 11(2), 127–138.

Graumann, M., Ciuffi, C., Dwivedi, K., Roig, G., & Cichy, R. M. (2022). The spatiotemporal neural dynamics of object location representations in the human brain. Nature Human Behaviour, 6(6), 796–811.

Grootswagers, T., Wardle, S. G., & Carlson, T. A. (2017). Decoding dynamic brain patterns from evoked responses: A tutorial on multivariate pattern analysis applied to time series neuroimaging data. Journal of Cognitive Neuroscience, 29(4), 677–697.

Harel, A., Groen, I. I., Kravitz, D. J., Deouell, L. Y., & Baker, C. I. (2016). The temporal dynamics of scene processing: A multifaceted EEG investigation. Eneuro, 3(5).

Hogendoorn, H., & Burkitt, A. N. (2018). Predictive coding of visual object position ahead of moving objects revealed by time-resolved EEG decoding. Neuroimage, 171, 55–61.

Johnston, P., Robinson, J., Kokkinakis, A., Ridgeway, S., Simpson, M., Johnson, S., … & Young, A. W. (2017). Temporal and spatial localization of prediction-error signals in the visual brain. Biological Psychology, 125,45–57.

Kaiser, D. (2022). Characterizing dynamic neural representations of scene attractiveness. Journal of Cognitive Neuroscience, 34(10), 1988–1997.

Kaiser, D., Azzalini, D. C., & Peelen, M. V. (2016b). Shape-independent object category responses revealed by MEG and fMRI decoding. Journal of Neurophysiology, 115(4), 2246–2250.

Kaiser, D., Häberle, G., & Cichy, R. M. (2020). Cortical sensitivity to natural scene structure. Human Brain Mapping, 41(5), 1286–1295.

Kaiser, D., Inciuraite, G., & Cichy, R. M. (2020). Rapid contextualization of fragmented scene information in the human visual system. Neuroimage, 219,p 117045.

Kaiser, D., Oosterhof, N. N., & Peelen, M. V. (2016a). The neural dynamics of attentional selection in natural scenes. Journal of Neuroscience, 36(41), 10522–10528.

Kaiser, D., Turini, J., & Cichy, R. M. (2019). A neural mechanism for contextualizing fragmented inputs during naturalistic vision. Elife, 8,p e48182.

Klein, L.K., Maeillo, G., Paulun, V. & Fleming, R.W. (2020). Predicting precision grasp locations on three-dimensional objects, PLOS Computational Biology, 16(8), e1008081.

Malik, A., Doerschner, K., & Boyaci, H. (in press). Unmet Expectations About Material Properties Delay Perceptual Decisions. Vision Research.

Oostenveld, R., Fries, P., Maris, E., & Schoffelen, J. M. (2011). FieldTrip: open source software for advanced analysis of MEG, EEG, and invasive electrophysiological data. Computational Intelligence and Neuroscience, 2011, 1–9.

Oosterhof, N. N., Connolly, A. C., & Haxby, J. V. (2016). CoSMoMVPA: multi-modal multivariate pattern analysis of neuroimaging data in Matlab/GNU Octave. Frontiers in Neuroinformatics, 10,27.

Paulun, V. C., Schmidt, F., van Assen, J. J. R., & Fleming, R. W. (2017). Shape, motion, and optical cues to stiffness of elastic objects. Journal of Vision, 17(1), 20–20.

Rao, R. P., & Ballard, D. H. (1999). Predictive coding in the visual cortex: a functional interpretation of some extra-classical receptive-field effects. Nature Neuroscience, 2(1), 79–87.

Robinson, J. E., Woods, W., Leung, S., Kaufman, J., Breakspear, M., Young, A. W., & Johnston, P. J. (2020). Prediction-error signals to violated expectations about person identity and head orientation are doubly-dissociated across dorsal and ventral visual stream regions. Neuroimage, 206,p 116325.

Schmid, A., Doerschner, K. (2018). The contribution of optical and mechanical properties to the perception of soft and hard breaking materials. Journal of Vision, 18(1):14, 1–32.

Schmid, A.C., Barla, P. & Doerschner, K. (preprint). Material category determined by specular reflection structure mediates the processing of image features for perceived gloss. bioRxiv, doi: https://doi.org/10.1101/2019.12.31.892083

Schmidt, F., Paulun, V. C., van Assen, J. J. R., & Fleming, R. W. (2017). Inferring the stiffness of unfamiliar objects from optical, shape and motion cues. Journal of Vision, 17(3): 18,1–17.

Sharan, L., Rosenholtz, R., & Adelson, E. H. (2014). Accuracy and speed of material categorization in real-world images. Journal of Vision 14(9), 12.

Stefanics, G., Astikainen, P., & Czigler, I. (2015). Visual mismatch negativity (vMMN): a prediction error signal in the visual modality. Frontiers in human neuroscience, 8,1074.

Tang, M. F., Smout, C. A., Arabzadeh, E., & Mattingley, J. B. (2018). Prediction error and repetition suppression have distinct effects on neural representations of visual information. Elife, 7,e33123.

Urgen, B. M., & Boyaci, H. (2021). Unmet expectations delay sensory processes. Vision Research, 181, 1–9.

Van Assen, J.J. Barla, P. & Fleming, R.W. (2018). Visual features in the perception of liquids. Current Biology, 28(3), 452–458.

van Driel, J., Olivers, C. N., & Fahrenfort, J. J. (2021). High-pass filtering artifacts in multivariate classification of neural time series data. Journal of Neuroscience Methods, 352,p 109080.

VanRullen, R. (2011). Four common conceptual fallacies in mapping the time course of recognition. Frontiers in Psychology, 2,p 365.

Wiebel, C. B., Valsecchi, M., & Gegenfurtner, K. R. (2014). Early differential processing of material images: Evidence from ERP classification. Journal of Vision, 14(7), 10–10.

